# Temperature affects viral kinetics and vectorial capacity of *Aedes aegypti* mosquitoes co-infected with Mayaro and Dengue viruses

**DOI:** 10.1101/2023.05.17.541186

**Authors:** Gerard Terradas, Jaime Manzano-Alvarez, Chiara Vanalli, Kristine Werling, Isabella M Cattadori, Jason L Rasgon

## Abstract

Increasing global temperatures and unpredictable climatic extremes have contributed to the spread of vector-borne diseases. The mosquito *Aedes aegypti* is the main vector of multiple arboviruses that negatively impact human health, mostly in low socioeconomic areas of the world. Co-circulation and co-infection of these viruses in humans have been increasingly reported; however, how vectors contribute to this alarming trend remains unclear. Here, we examine single and co-infection of Mayaro virus (-D strain, *Alphavirus*) and dengue virus (serotype 2, *Flavivirus*) in *Ae. aegypti* adults and cell lines at two constant temperatures, moderate (27°C) and hot (32°C), to quantify vector competence and the effect of temperature on infection, dissemination and transmission, including on the degree of interaction between the two viruses. Both viruses were primarily affected by temperature but there was a partial interaction with co-infection. Dengue virus quickly replicates in adult mosquitoes, with a tendency for higher titers in co-infected mosquitoes at both temperatures and mosquito mortality was more severe at higher temperatures in all conditions. For dengue, and to a lesser extent Mayaro, vector competence and vectorial capacity were higher at hotter temperature in co- vs single infections and was more evident at earlier timepoints (7 vs 14 days post infection). The temperature-dependent phenotype was confirmed *in vitro* by faster cellular infection and initial replication at higher temperatures for dengue but not for Mayaro virus. Our study suggests that contrasting kinetics of the two viruses could be related to their intrinsic thermal requirements, where alphaviruses thrive better at lower temperatures compared to flaviviruses, but further studies are necessary to clarify the role of co-infection at different and variable temperature regimes.

**Author summary:** Global warming is having devastating consequences for the environment, and a cause of concern is the increase in local abundance and geographic range of mosquitoes and the associated viruses they transmit. This study explores how temperature affects the mosquito’s ability to survive and potentially spread two viruses, Mayaro and dengue, in single or co-infections. We found that Mayaro virus was not clearly affected by temperature or the presence of dengue infection. In contrast, dengue virus showed higher infection and potential for higher transmission in mosquitoes kept at high temperatures, and this trend was stronger in co-infections compared to single infections. Mosquito survival consistently decreased at high temperatures. We hypothesize the differences observed for dengue virus are due to the faster growth and viral activity in the mosquito at hotter temperatures, a pattern not observed for Mayaro virus. More studies under different temperature regimes are needed to clarify the role of co-infection.

## Introduction

The mosquito *Aedes aegypti* is broadly distributed throughout the tropics/sub-tropics (1) and is the vector of multiple arboviruses such as dengue (DENV), Chikungunya (CHIKV) and Zika (ZIKV). More recently, *Ae. aegypti* has also been reported to be a competent vector for Mayaro virus (MAYV), an emerging alphavirus originally isolated in 1954 in Trinidad and Tobago and which is currently present in many regions of Central and South America (2, 3). In endemic areas that harbor extensive mosquito populations, arboviruses frequently co-circulate (4) with some evidence of synchronous outbreaks in human populations (5) and local hotspots of infection (6, 7). The endemic co-circulation of these viruses, and the expansion of their vectors into new geographical areas has been frequently associated with climate change and human development (8). The increasing trend of human cases in the last decades has been a serious public health concern, aggravated by the lack of commercially available therapeutics and successful prophylactics, including the ability to provide effective diagnostics.

The ecology of *Ae. aegypti* and viral kinetics are strongly regulated by temperature. Non-linear thermal responses have been described for many mosquito traits, such as larval development, adult survival, infection probability and, for virus-mosquito interactions, the extrinsic incubation period (EIP) (9, 10). Simulations based on thermal-dependent traits suggest that *Ae. aegypti* optimal temperature for viral transmission peaks at 29.1°C while no transmission occurs outside the range between 17.8°C and 34.6°C under regimes of constant temperature (10). This, coupled with the fact that DENV transmission peaks at temperatures over 30°C (11, 12) and CHIKV (and likely other alphaviruses) is less active at those temperatures (13), raises the question of whether the interactions between viruses, and their co-transmission, could be affected by thermal variations. For instance, viruses have different thermal requirements that can impact how they interact at different temperatures (reviewed extensively in (14)), while mosquito thermal tolerance could disproportionately hamper their ability to endure multiple viral infections with consequences for their fitness and virus transmission (15). Importantly, given the strong non-linearities of thermal responses of both mosquitos and viruses, it is difficult to predict the outcomes of co-infections based on our knowledge of virus-mosquito interactions from single infections (16). Therefore, in addition to the intrinsic properties of *Ae. aegypti* and its arboviruses, we need to consider the effect of temperature on each player and their relationships when investigating infection patterns.

With the spread of arboviruses (8, 17) and the increase in human cases, there have been a growing number of patients presenting co-infection combinations of CHIKV, DENV and ZIKV (18) and recently, reports of MAYV with DENV (19), CHIKV (20, 21) and ZIKV (22). While the observations of multiple infections in same patients indicate that *Ae. aegypti* co-infection is possible and likely more widespread than expected, whether these co-infections are the result of sequential bites from single-infected mosquitoes or the result of a single bite from a mosquito infected with multiple viruses is still fundamentally unknown. Moreover, local investigations both on humans and vectors are lacking (but see (23, 24)), making it difficult to generalize trends observed in humans to *Ae. aegypti* populations.

The competence of *Ae. aegypti* to single and concurrent infections with CHIKV, DENV, ZIKV and more recently MAYV has been documented. In laboratory experiments, *Ae. aegypti* has been found to be able to concurrently carry and simultaneously transmit combinations of these four arboviruses (25-30). Several studies found that there can be some level of interference based on virus pairing that affects total viral load and capacity of dissemination or transmission when compared to single infections (25, 27, 31). Other studies found no clear evidence of interaction (25, 28, 29, 32) or refractoriness in the case of co-infections of DENV and CHIKV in *Ae. aegypti* (33)). The general observation from available studies suggests that there is no consistency in the infection outcomes and some of these trends also appear to be affected by the order of infection, namely, whether there is sequential or simultaneous intake. For instance, CHIKV-DENV co-transmission was low in *Ae. aegypti* sequential infections but null when simultaneously dosed (27). Similarly, the sequential infection of MAYV-ZIKV produced different outcomes based on which infection occurred first and mostly affecting the latter one (26). While differences in virus strains and mosquito lines are important contributors of these results, laboratory settings, like different temperature conditions, could also play a role.

In this study, we focused on the interactions between DENV and MAYV, since both are endemic in Central and South America and the likelihood of co-infection in *Ae. aegypti* is increasing. Indeed, although MAYV outbreaks in humans have been associated with the vector *Haemagogus janthinomys*, there are reports of natural *Ae. aegypti* infections, which could increase the risk of potential urban cycles for MAYV and co-circulation with the already established DENV (34). Currently, MAYV infection has been reported in a wide range of animal species with seroprevalence ranging between 21% and 72% (2). In tropical regions, similar transmission abilities have been reported between MAYV and other arboviruses (basic reproduction number R_0_ range: MAYV= 1.1–3.5, DENV= 4.25, ZIKV=-2.98 and CHIKV= 3.09) (2). However, vectorial capacity might change if the prevalence of co-infected mosquitoes increases, or warmer temperatures unevenly affect areas suitable to *Ae. aegypti* (35, 36).

Here, we examined how temperature impacts vector infection, survival and viral kinetics in laboratory experiments of *Ae. aegypti* infected with either DENV or MAYV, or simultaneously, under two constant temperatures, 27°C (moderate) and 32°C (high). A complementing *in vitro* study was performed to provide additional insight on virus interaction within cells and the effect of temperature on viral growth over time. Finally, using our laboratory data and available literature, we estimated vector competence and vectorial capacity in single and co-infections under different temperatures to gain better understanding of thermal dependencies of *Ae. aegypti* co-infections and differences from single infections.

## Methods

### Mosquitoes

#### Aedes aegypti

Liverpool strain mosquito eggs (NR-48921) were provided by BEI Resources (Manassas, VA). Insects were maintained and reared at the Millennium Sciences Complex insectary (The Pennsylvania State University, University Park, USA) at 27°C, 12:12h light:dark cycle and 80% humidity. Larvae were fed fish food pellets (Tetra, Germany) and adult mosquitoes were kept in 30×30×30cm cages on 10% sucrose diet *ad libitum*. For colony breeding and maintenance, adult females were allowed to feed on anonymous human blood using membrane glass feeders following a previously described protocol (37).

### Cells

Three types of cells were used in our study: African green monkey kidney cells (Vero; CCL-81) were cultured at 37°C and 5% CO2 in complete media consisting of Dulbecco’s modified Eagle’s medium complemented with 10% fetal bovine serum (FBS) and 1% penicillin/streptomycin (PenStrep). *Aedes albopictus* RNA interference-deficient cells (C6/36; CRL-1660) were kept and passaged at 28°C in RPMI media supplemented with 10% FBS and 1% PenStrep. For each passage, both Vero and C6/36 cells were detached by trypsinization (0.25% trypsin; Corning Inc., Corning, NY) and diluted in their respective fresh complete media. *Ae. aegypti* cells (Aag2; kind gift from Elizabeth McGraw, The Pennsylvania State University) were grown and passaged in Schneider’s insect medium supplemented with 10% FBS and 1% PenStrep. All cell culture reagents were purchased from Gibco, Thermo Fisher Scientific (Waltham, MA).

### Viruses

We used MAYV strain BeAn343102 (BEI Resources, Manassas, VA), corresponding to a genotype D strain (MAYV-D) isolated in May 1978 from a monkey in Para, Brazil. The DENV-2 strain (herein referred to as DENV) was isolated from a human patient in Timor-Leste in 2000 (ET300; GenBank accession number EF440433.1). MAYV stocks were grown in Vero cells (at 37°C), whereas DENV was passaged in C6/36 cells (at 28°C) until supernatant collection. Viruses were allowed to infect cells at a multiplicity of infection (MOI) of 0.1 for an hour, viral inoculum was then removed and replaced with fresh media containing 2% FBS. Virus-infected supernatant was aliquoted at 24 hpi for MAYV (corresponding to the time when a clear cytopathic effect on the cells was observed) and 7 dpi for DENV, then stored at −80°C until use for mosquito infections and viral titration. Viral stock titers were obtained using FFAs (ffu/mL), as described below.

### Temperature-dependent single and co-infection *in vivo*

Mosquitoes were reared as previously stated until emergence. At that point, they were split in different cages and transferred to 27°C (temperate) and 32°C (hot) incubators, both kept at standard 80% humidity. After 7 days in their respective temperature treatments, mosquitoes were fed virus-spiked blood containing 5×10^6^ ffu/mL per virus (i.e. either MAYV or DENV, or both) and those engorged were sorted, split in independent cups per type of infection (single or co-infection) and moved back to each experimental temperature (27°C or 32°C) treatment. Whole mosquito bodies were collected immediately after blood feeding to ensure that the uptake of infectious virus occurred at similar concentrations and that the delivered virus was infective (S1 Fig.). While in incubators, mosquito mortality for each condition was assessed daily. At 7- and 14-days post infection (dpi), mosquitoes were anesthetized using triethylamine (Sigma-Aldrich) and then individually forced to salivate to assess their viral transmission capacity. After 30 minutes of forced salivation, each female’s midgut was dissected to assess the level of mosquito infection, while viral presence in the carcass (rest of the body) was used as a proxy for dissemination to body organs. Tissue samples were homogenized using a TissueLyser II (QIAGEN, Germany) and all samples were stored at -80°C until titration. Fluorescence-based focus-forming assays (FFAs) were used to visually count the presence of infectious viral particles in each sample.

### Temperature-dependent single and co-infection *in vitro*

We performed *in vitro* experiments using a well-established *Ae. aegypti* cell line (Aag2), which is typically kept and grown at 27°C. We adapted the line to hotter conditions (32°C) to perform the experiments at the same temperatures used for the *in vivo* experiments. Cells were seeded at 80-90% confluence in 12-well plates and, the following day, infected as mock, single or co-infections of MAYV and DENV at a multiplicity of infection (MOI) of 0.1. After 1h of incubation at 37°C, the viral inoculum was removed, cells washed twice to remove unbound virus, and fresh medium added. Viral supernatants were collected at selected timepoints for 3 days, in intervals of 6h to 12h. At each time point, 100μL of virus-containing supernatant was collected and stored at -80°C. Upon sampling, 100uL of fresh media were added to each well to maintain the same initial volume. Three technical replicates were performed for each condition, and the experiment was run twice. Supernatant samples were tested for positive viral contents using FFAs and human Vero cells in the same fashion as adult mosquito samples.

### Fluorescence-based Focus-forming assay (FFA)

Vero cells were counted using a hemacytometer and plated in DMEM complete media the day before infection to obtain 80-90% confluency in 96-well plates. The next day, media was removed and cells were incubated with 30 µL of 10-fold dilutions of each homogenized tissue sample in FBS-free media. Saliva samples were not diluted due to their lower titers. After an hour of incubation, the viral media was removed and cells were kept on an overlay of 0.8% methylcellulose (or CMC) in complete media for 24h. After fixation and permeabilization of the cells, and extensive PBS washes, viral antigens were labeled using specific monoclonal antibodies, specifically, mouse monoclonal anti-CHIKV E2 envelope glycoprotein clone CHK-48 (α-CHK-48, BEI Resources) for MAYV and mouse monoclonal anti-flavivirus clone D1-4G2-4-15 (BEI Resources) for DENV, both diluted 1:500 in PBS. The next day, cells were washed thoroughly with cold PBS to remove unbound primary antibody and a secondary antibody (Alexa 488 goat anti-mouse IgG secondary antibody (Invitrogen, OR Waltham, MA)) at a 1:750 dilution in PBS was applied to the samples for 1 hour at room temperature. Cells were rinsed of antibodies prior to viral evaluation. Green fluorescence was observed using a fluorescein isothiocyanate (FITC) filter on an Olympus BX41 microscope with a UPlanFI 4× objective. Foci were counted manually in the appropriate dilution, and the viral titers were back calculated to ffu/mL.

### Statistical analysis

We used a zero-inflated negative binomial regression (ZINBR) model to examine differences in MAYV or DENV titers between temperatures (27°C or 32°C), treatments (single or co-infection), and their pairwise interaction, for midgut and carcass (which account for viral infection and dissemination within the mosquito, respectively) at two timepoints, 7 and 14 days post-infection, examined independently. For saliva samples (indicatives of potential transmission) we performed a negative binomial generalized linear model (GLM-NB) as the ZINBR could not be applied because of the lack of non-zero data in some of the groups. To test for daily differences in adult mosquito mortality by temperature and treatment, we used survival log-rank Mantel-Cox test. To investigate whether temperature and co-infection play a role in *in vitro* intracellular viral kinetics, linear regression of log-transformed FFA data by time were estimated for every treatment and temperature, independently, and the related slopes and intercepts were then compared.

### Temperature-dependent vector competence and vectorial capacity in single and co-infection

Experimental data and data from the literature were used to estimate vector competence and vectorial capacity for MAYV and DENV in single and co-infection and each constant temperature (27°C and 32°C). The vectorial capacity *V* describes the potential of a vector to transmit a pathogen and is defined as the number of new mosquito infectious bites that arise from one infected person introduced into a population of entirely susceptible hosts in a day (eq. 1).

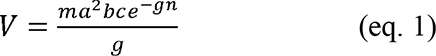

Here, *m* is the mosquito to human ratio, *a* is the mosquito biting rate, *b* is the mosquito to human probability of transmission and *c* is the human to mosquito transmission probability, while *g* is the mosquito mortality rate and *n* represents the extrinsic incubation period (EIP) (Table 2). This latter parameter is defined as the duration of the period from ingestion of the infective blood meal until the mosquito becomes infectious in the salivary glands. Under this assumption, the probability of mosquito survival during this period is *exp(-gn)*. We estimated the mosquito mortality rate as the reciprocal of the median survival time in days for each temperature setting and treatment. Vector competence is defined as the probability of transmitting the pathogen given host exposure and is calculated as the product between the parameters *b* and *c*. The parameters *b*, *c* and *g* can be estimated from our experiments, while we performed a literature search to gather plausible values for the remaining parameters (*m*, *a, n*). Here, we assumed an equal proportion of mosquito per human (*m*=1), a common assumption for vector-borne disease models of transmission (38). We found evidence of a temperature dependence of *Ae. aegypti* biting behavior (39, 40) which we assumed is independent from treatment. We also gathered information on the temperature dependence of DENV’s EIP (41) and, since there is little information for MAYV, we extrapolated this information from CHIKV as it is the closest representative with temperature dependencies available (42). For high temperature settings the shortest reported EIP is 2 days (43), while as a proxy of moderate temperature we selected the EIP of 7.15 days (i.e. average value calculated from (44)) and we used the former value to represent EIP at 32°C and the latter at 27°C.

## Results

### Temperature unevenly affects DENV and MAYV in single and co-infections

After 7 days in their respective temperature treatments, adult *Ae. aegypti* females were orally challenged with a blood meal spiked with 5×10^6^ ffu/mL of either MAYV (single), DENV (single), or both (for co-infection, total of 1×10^7^ ffu/mL). Tissues (midgut, carcass (rest of the body) and saliva) of female mosquitoes that took an infectious blood meal were collected at 7 and 14dpi to assess vector competence with viral titers and prevalence of positive mosquitoes.

Overall, MAYV and DENV exhibit different kinetics by temperature and treatment (Fig 1, Table 1). For MAYV, midgut infection titers are negatively associated with temperature at 14dpi, while no significant relationships are found with treatment at either collection timepoint (Fig 1a and 1b, Table 1). There is a significant negative effect of both temperature and treatment on MAYV carcass dissemination at 7dpi but not at 14dpi (Figure 1c and 1d, Table 1). For saliva, temperature positively affects MAYV titers at both 7dpi and 14 dpi (Figure 1e, 1f, Table 1) and titers significantly increase for co-compared to single infected mosquitoes at 14dpi (Fig 1e, Table 1). Regarding mosquito competence to MAYV, there is no significant difference in the prevalence of positive mosquitoes between treatment or temperature for infection, dissemination, or transmission at any of the timepoints (Fig 1g).

**Fig 1.**
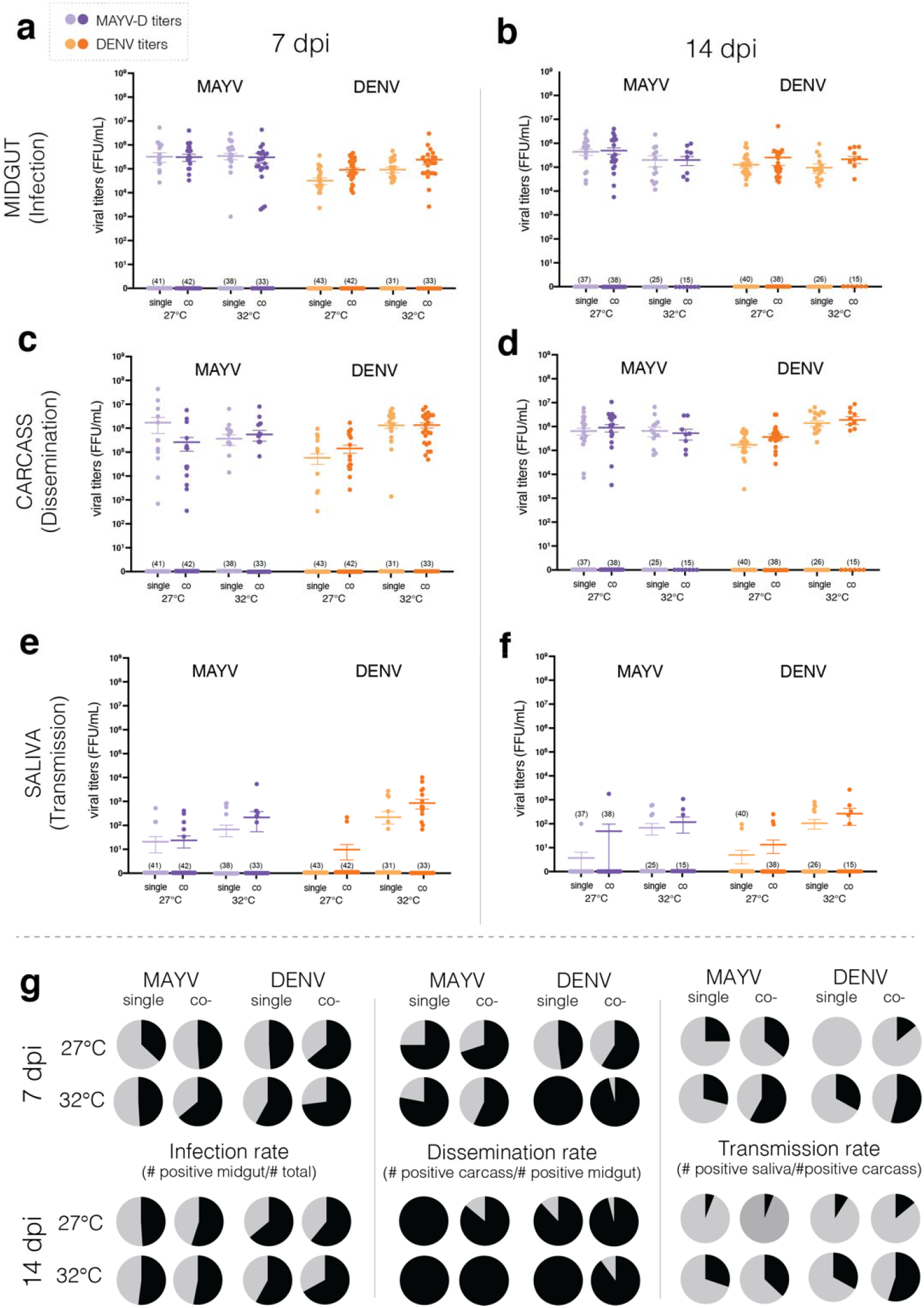
Mayaro (MAYV) and dengue (DENV) titers in adult *Ae. aegypti* tissues. Tissues were collected 7 and 14 days after bloodfeeding with a single or a co-infection dose of MAYV and DENV. Viral titers for infection (**a, b** - midgut), dissemination (**c, d** - carcass) and transmission (**e, f** - saliva) were assessed using FFA, and plotted based on virus and time after challenge. Each point represents a sample of an individual mosquito tissue. **(g)** Mosquito infection, dissemination and transmission rates by temperature and treatment at 7 and 14 dpi are plotted in **a-to-f**. Positive % in black, negative % in gray.

**Table 1.**
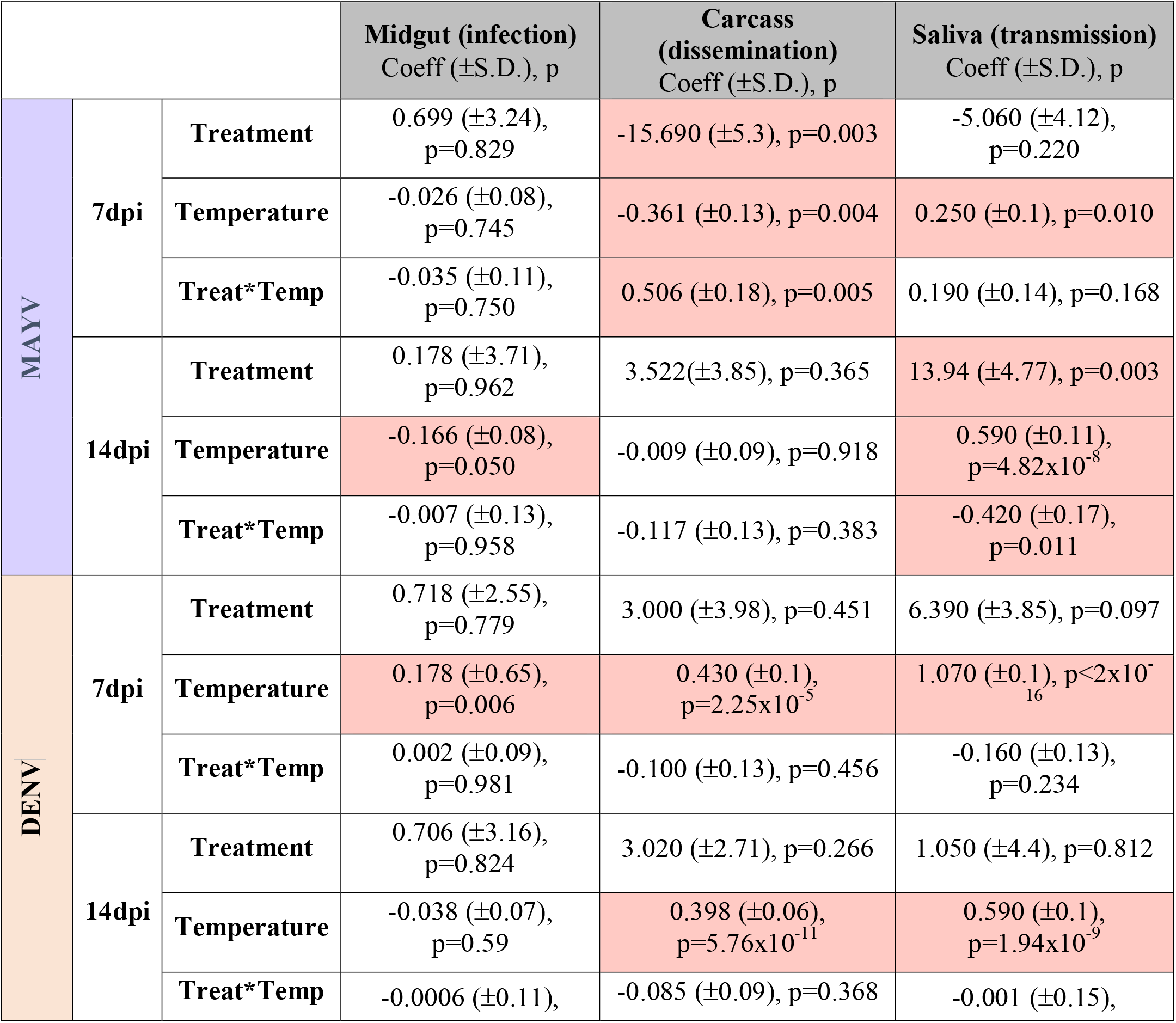
Summary of Zero-Inflated (midgut, carcass) and Generalized Linear (saliva) models. Estimated coefficients (± standard deviation) and p-values are reported. Significant effects (p<0.05) are shaded in red.

For DENV, titers are significantly lower at 27°C than 32°C for every assessed tissue at both 7dpi and 14 dpi (Figure 1a, 1c and 1e, Table 1), except for the midgut at 14dpi. A positive relationship with temperature is also found for saliva titers at both timepoints (Figure 1e and 1f, Table 1). However, no significant relationships are observed with treatment, as viral titers do not differ between single and co-infections. Differences in prevalence rates were examined using Fisher’s exact test depending on treatment and infection (Figure 1g). For those, we observed statistical differences in both dissemination and transmission rates depending on treatment at 14dpi (Fig 1g).

To test whether higher temperatures affect mosquito fitness when infected with one or both viruses, we performed survival analysis on infected *Ae. aegypti* over a 14-day period. In general, for all the infection groups, mosquitoes perish significantly faster at 32°C than 27°C (X^2^= 331.3, df=7, p<0.0001; 7dpi: mean percentage of mortality at 32°C=44% vs 27°C=7%; 14dpi: mean percentage of mortality at 32°C=73% vs 27°C=28%; Fig 2a). Pairwise comparison of each viral treatment by temperature, independently, shows a significant reduction of survival at 32°C for all the cases (Uninfected: X^2^= 13.1, df=1, p=0.0003; MAYV: X^2^= 89, df=1, p<0.0001; DENV: X^2^= 47.4, df=1, p<0.0001: MAYV-DENV: X^2^= 75.3, df=1, p<0.0001; Figure 2b). Similarly, when we compare different treatments within the same temperature, uninfected mosquitoes or those exposed to DENV only are the least impacted and significantly differ from the groups challenged with MAYV, which show the highest mortality (27°C: X^2^=13.1, df=3, p=0.0044; 32°C: X^2^=53, df=3, p<0.0001). Likewise, when comparing MAYV-challenged mosquitoes, there is a tendency for higher mortality of the co-infected group at 27°C (X^2^=3.11, df=1, p=0.077), which appears to increase at later timepoints (i.e. 11dpi onward), but no differences are found at 32°C (X^2^=0.13, df=1, p=0.723; Fig 2a).

**Fig 2.**
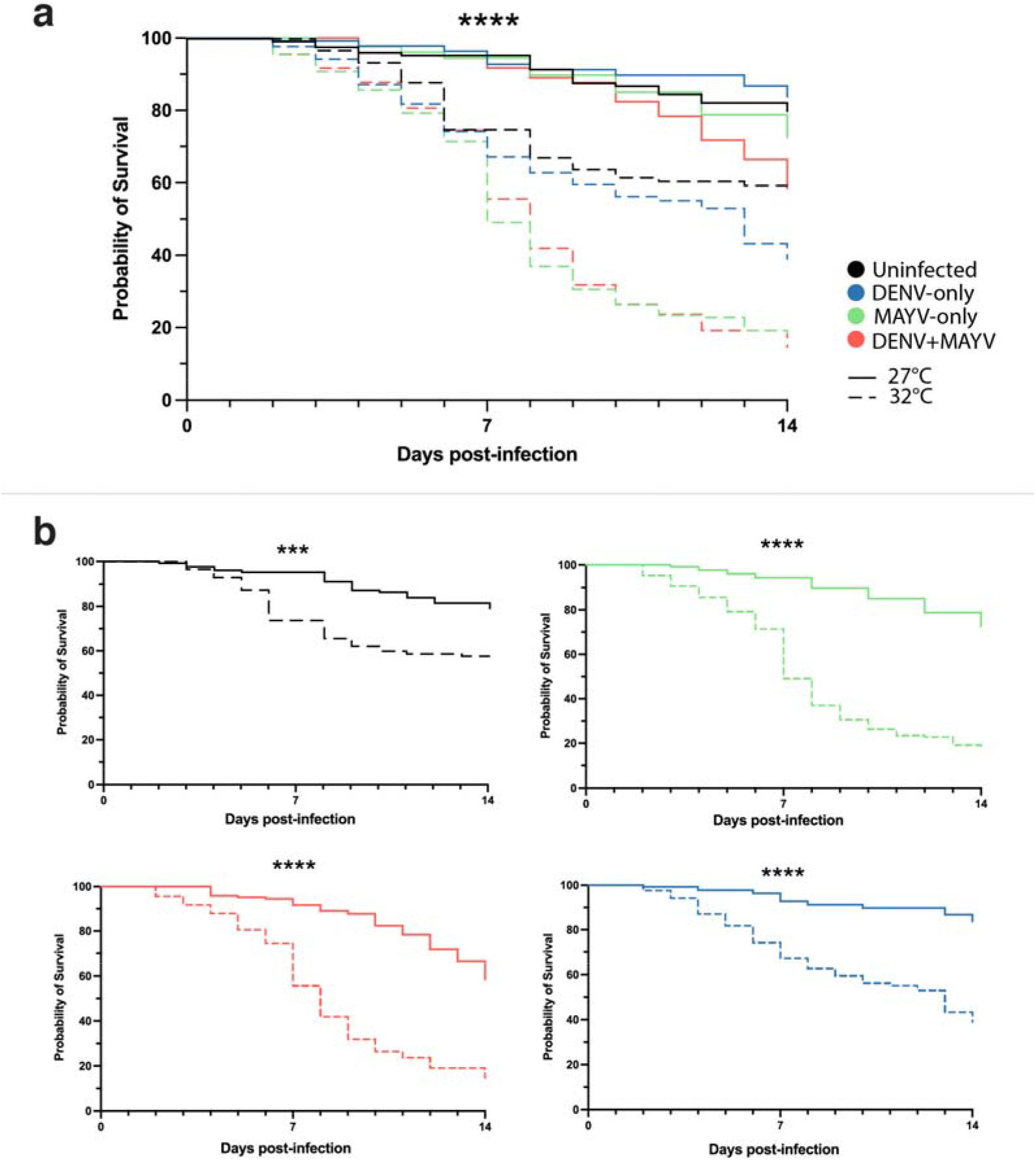
Mosquito survival curves after viral challenge. Dead mosquitoes were counted daily and graphed **a)** altogether over time or **b)** separated by treatment type. Significant result as follows: *** p<0.001, **** p<0.0001.

### Temperature affects single and co-infection kinetics *in vitro* in a virus-specific manner

To examine whether temperature and treatment have an effect on intracellular viral kinetics, viral growth curves were assessed using the well-established *Ae. aegypti* cell line (Aag2) (Fig 3a). Curves were compared by their slopes, which indicate how fast cellular replication occurs, and intercepts, which explains how early growth starts. No significant differences are found for MAYV replication between treatments or temperatures (slopes: F=0.297, df=3, p=0.827 and intercepts: F=1.625, df=3, p=0.185) (Fig 3b). In contrast, DENV shows significant differences in the initial conditions (intercepts: F=18.95, df=3, p<0.0001) but not in the intracellular growth (slopes: F=1.095, df=3, p=0.353) (Fig 3c). These results suggest that co-infection and/or warmer temperatures appear to have no effect on intracellular replication, but both conditions promote faster DENV replication at the early stages of the infection.

**Fig 3.**
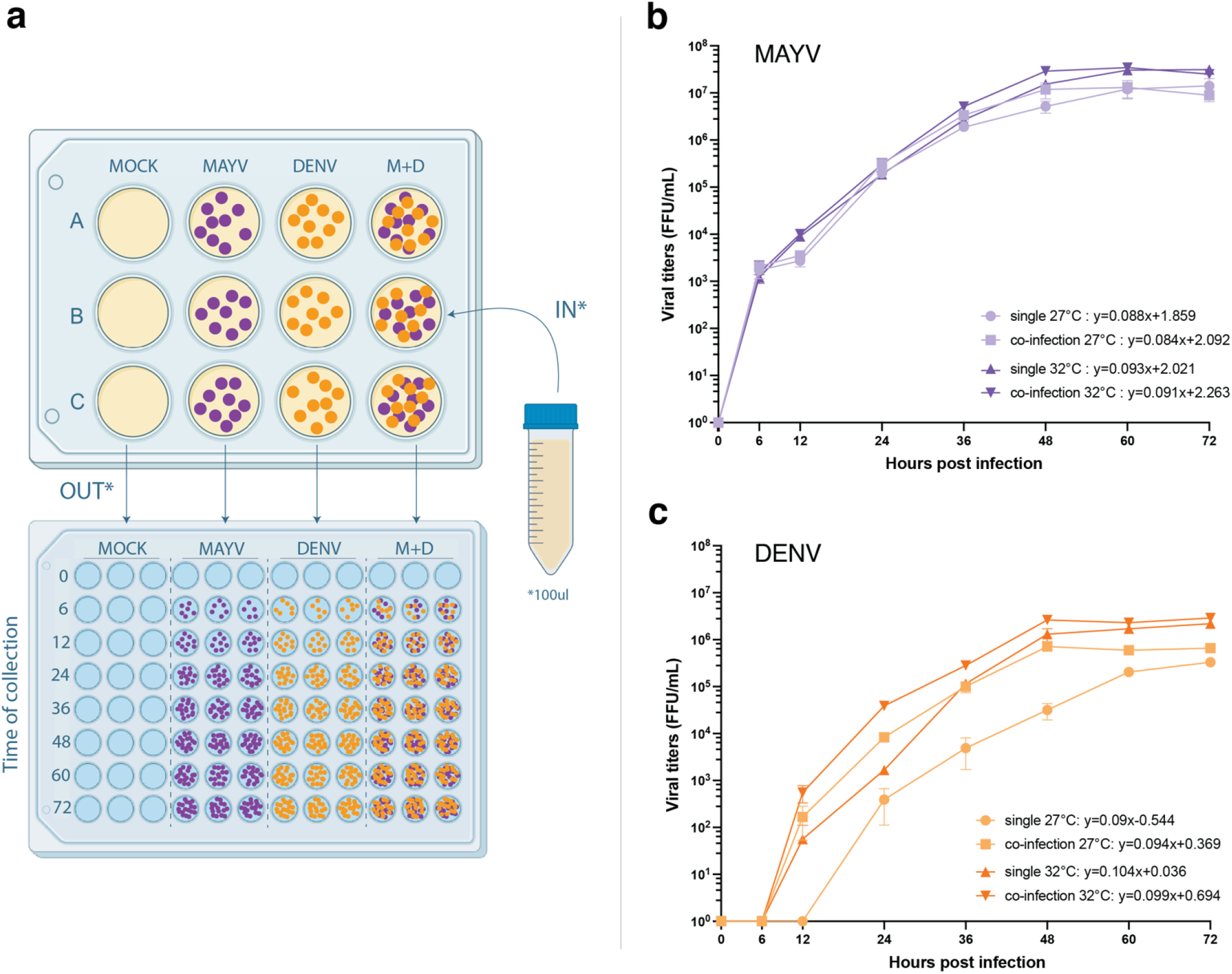
MAYV and DENV growth curves in Aag2 cells. Experimental schematics and collection processes. Viral titers in cell supernatants were collected at fixed time points post infection to assay viral growth at 27°C and 32°C for single and co-infection treatments, trials were run twice and each including three technical replicates **(a)**. Viral titers for MAYV **(b)** and DENV **(c)** were analyzed using FFA and plotted as log-mean (+-SE) by time post infection for each virus and treatment.

### Temperature affects vector competence and vectorial capacity in single and co-infection

Three parameters are estimated from the current study: mosquito mortality, mosquito-to-human transmission and human-to-mosquito transmission. Confirming the statistical results, mosquito mortality rate, *g*, is consistently higher at 32**°**C than at 27**°**C, but there is a possible co-infection effect where mortality rate almost doubles for mosquitoes with DENV-MAYV compared to the DENV-only group; this trend is lacking for MAYV (Table 2). The mosquito-to-human probability of transmission, *b*, is higher at warmer temperatures at both 7 and 14dpi, and the presence of the second virus contributes to the increase of the transmission probability for both DENV and MAYV (Table 2). This same pattern is also found for the human-to-mosquito probability of transmission, *c*, as co-infected mosquitoes have higher competence than single-infected ones, although this trend is less consistent at 14dpi. Here, we assumed that biting rate was temperature-dependent but not related to treatment, and we show that higher biting rate is associated with warmer temperatures, consistent with previous studies.

**Table 2.**
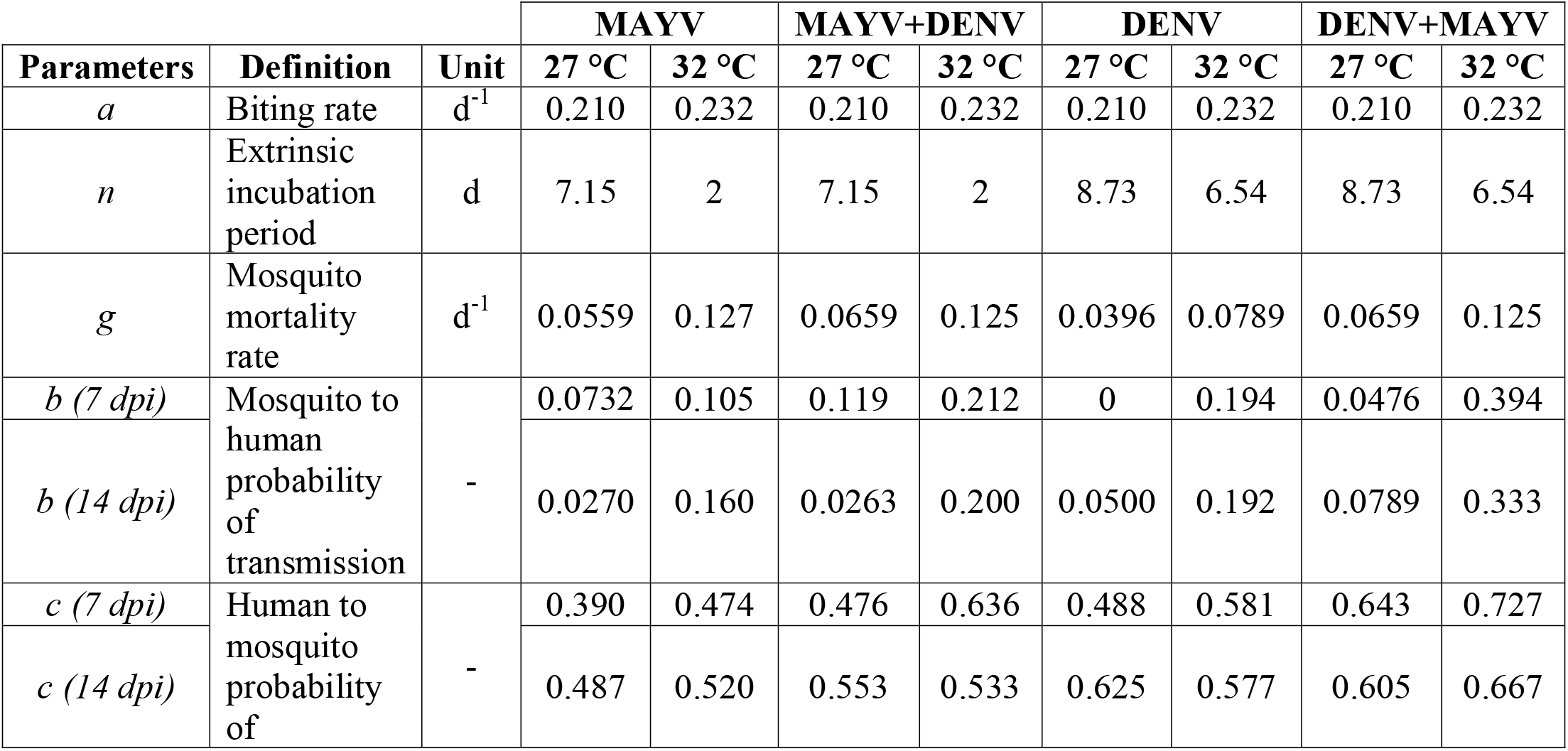
Parameter definitions, unit of measure and relative values at 27**°**C and 32**°**C and at 7- and 14-days post infection (dpi) of MAYV and DENV in single and co-infection.

The described parameters are then used to evaluate thermal differences in vector competence and vectorial capacity for MAYV and DENV in single- and co-infected mosquitoes. For both viruses, we show an increase in *Ae. aegypti* competence with temperature from 27°C to 32°C (Figure 4a and 4b). The highest increase is found for DENV in co-infected mosquitoes (at 7 and 14dpi, respectively: +0.256 and +0.174) and the lowest for MAYV in single-infected vectors (at 7dpi: +0.021). For both viruses, differences in *Ae. aegypti* competence between single- and co-infection are greater at 32**°**C, suggesting an interaction between coinfection and temperature in determining the probability of virus transmission. The vectorial capacity for MAYV and DENV appears to be weakly related to temperature or co-infection, probably because mosquito survival is lower at warmer temperature and when carrying both viruses, despite more efficient transmission (Figure 4c and 4d, Table 2). Specifically, DENV vectorial capacity at 32**°**C exhibits low variability across infection types. For MAYV, vectorial capacity shows a similar trend for single and dual-infected mosquitoes, albeit with more variability and with a negligible relationship with temperature for single-infected mosquitoes at 7dpi (+0.00124). Overall and for both viruses, vectorial capacity is at the highest in co-infected mosquitoes at 32**°**C and at 7dpi.

**Fig 4.**
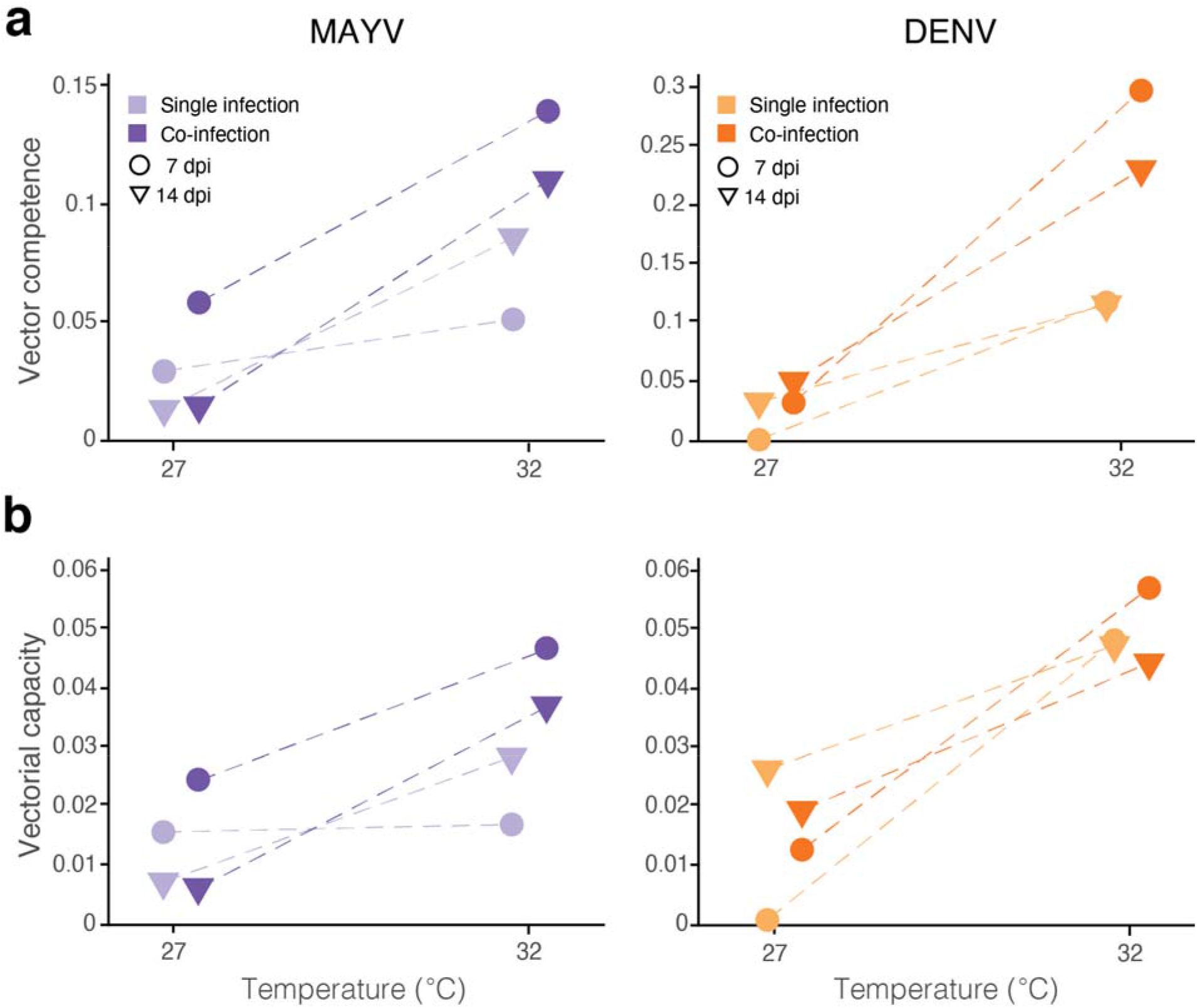
**(a)** Vector competence and **(b)** vectorial capacity of MAYV (left panels) and DENV (right panels) in single infection (lighter shades) and co-infection (darker shades) by temperature at 7dpi (circles) and 14dpi (triangles). Dashed lines are provided for tracking the same infection.

## Discussion

Thermal dependencies of *Ae. aegypti* and the common arboviruses they transmit have been predominantly studied in the laboratory as single infections while the role of temperature on viral co-infection in mosquitoes has been poorly investigated. This line of research is important not only for advancing the understanding of the epidemiology of vector-borne infections but also to appreciate the impact of climate warming on the fitness of mosquito vectors and the transmission of co-circulating viruses. We examined the impact of moderate and hot temperatures on the kinetics of MAYV and DENV in *Ae. aegypti* with single or dual infections and show that 1) both viruses can be co-transmitted by adult mosquitoes and the presence of the second virus appears to partially affect both MAYV and DENV kinetics, 2) warmer temperature contributes to higher replication and infectivity, and mosquito mortality, especially for DENV and 3) estimation of vector competence and vectorial capacity support the important role of co-infection, particularly at higher temperatures.

We found similar rates of infection (positive titers in midgut) of MAYV and DENV at all conditions and timepoints, with a partial effect of temperature but not treatment, suggesting that the capacity of the viruses to successfully infect the mosquito occurs irrespective of these conditions. We however observe increased dissemination rates and titers for DENV at 32**°**C at 7dpi, and these same quantities (rate and titer) are increased for transmission of both MAYV and DENV at early and late timepoints, irrespective of infection type (Fig 1g). This is relevant in the context of global warming, which not only affects the distribution of mosquitoes and viruses they transmit, but also the suitability of DENV transmission in endemic areas affected by an increase in temperature (12). In essence, the contribution of higher temperature, first, and co-infections, second, is less clear for the infection success of MAYV than DENV, because at 27°C MAYV is already close to its optimal infecting temperature despite the slightly more rapid replication observed at 32°C. Our results are in accordance with general findings that higher temperatures are optimal for flaviviruses (10, 11) as they facilitate replication, body tissue dissemination, and transmission. This contrast with alphaviruses as they have more successful replication at moderate temperatures (13, 45). As such, the increase of temperature does not affect MAYV infection and replication nearly as much as it does for DENV, which infection in the mosquito appears to be more strongly driven by temperature as higher temperatures lead to higher viral titers in all tissues. Flavivirus infections have been observed to be negatively affected by the presence of MAYV (26) or a different alphavirus (CHIKV (25), Sindbis virus (46)). This general trend has been reported in many other studies (25, 26, 29, 46) and consistently show that alphaviruses can negatively impact the ability of flaviviruses to infect and disseminate although there was no effect on transmission. These differences could be associated to intrinsic behavioral singularities between viruses of the same genus as well as genotype x genotype (GxG) interactions where different strains, origin or genomic polymorphisms in specific mosquito and virus species affect their relationship (47, 48). Consistent with previous literature, we did not observe an effect of flavivirus on alphavirus infections.

The increase in rates and/or titers with temperature, albeit not always significant, may not be as worrisome as expected because these trends were also associated with higher mosquito mortality (Fig 2). This suggests that while mosquito are sensitive to temperature, this is exacerbated in infected mosquitos in a non-linear way, indicating they quickly lose fitness to thrive as temperature increases. Under temperature warming, multiple blood-feeding events would be less likely to occur, reducing the chances of an infected mosquito to encounter humans and transmit infections. However, when we consider the estimated vectorial capacity, the transmission potential of both MAYV and DENV are at the highest at 32°C at 7dpi for single and dual-infections, suggesting that the net effect of mosquito-virus interactions ultimately results in a higher risk of transmission as temperature increases. Indeed, while mosquito mortality rate *g* increases the probabilities of transmission, *b* and *c* increase faster, and are higher in dual than single infected mosquitos. Our findings suggest that MAYV-DENV infected mosquitoes could potentially contribute to higher virus co-circulation and, notably, human co-infection. However, more work in needed to disentangle how temperature changes and co-infections interact and affect vectorial capacity, for example the role of fluctuating temperature, and thus a more environmentally relevant pattern, both on vectors and viruses or the contribution of sequential versus concomitant infections.

Finally, our experiments assume that each virus contributes with a similar initial titer level and consistent in single and co-infections, a scenario that most likely does not reflect the more variable field conditions. In this respect, the inclusion of experiments testing different viral doses under different climatic scenarios can provide more natural conditions and increase biological accuracy. Moreover, it is also possible that mosquitoes acquire multiple viruses on separate blood-feed events (i.e. super-infection scenario) and thus the kinetics of the two viruses could be affected by the order of infection as reported by Brustolin et al. (26) where the sequence of MAYV-CHIKV super-infections was critical to flaviviral success. This is critical if we want to have a better understanding to the processes modulating vector co-infection and, ultimately, whether co-infected mosquitos play a key role to the spread and emergence of vector-borne infections. Our study sheds new light in this direction by providing fundamental knowledge in how temperature changes modulate mosquito-virus co-infections. As vectors are expected to expand their geographic range with climate warming, the interactions between mosquitoes and the pathogens they transmit are also likely to be affected by these changes; disentangling the processes behind these changes is a public health priority.

## Acknowledgements

We are grateful to Abby Greb from the Rasgon lab for helping with the mosquito mortality counting.

## Supporting information

Supporting Information files do not require full captions; only labels (“S1 Fig”) are fully required. Supporting Information files should be saved as “S1_Fig.tif”, “S1_File.pdf”, etc.

**S1 Fig.**
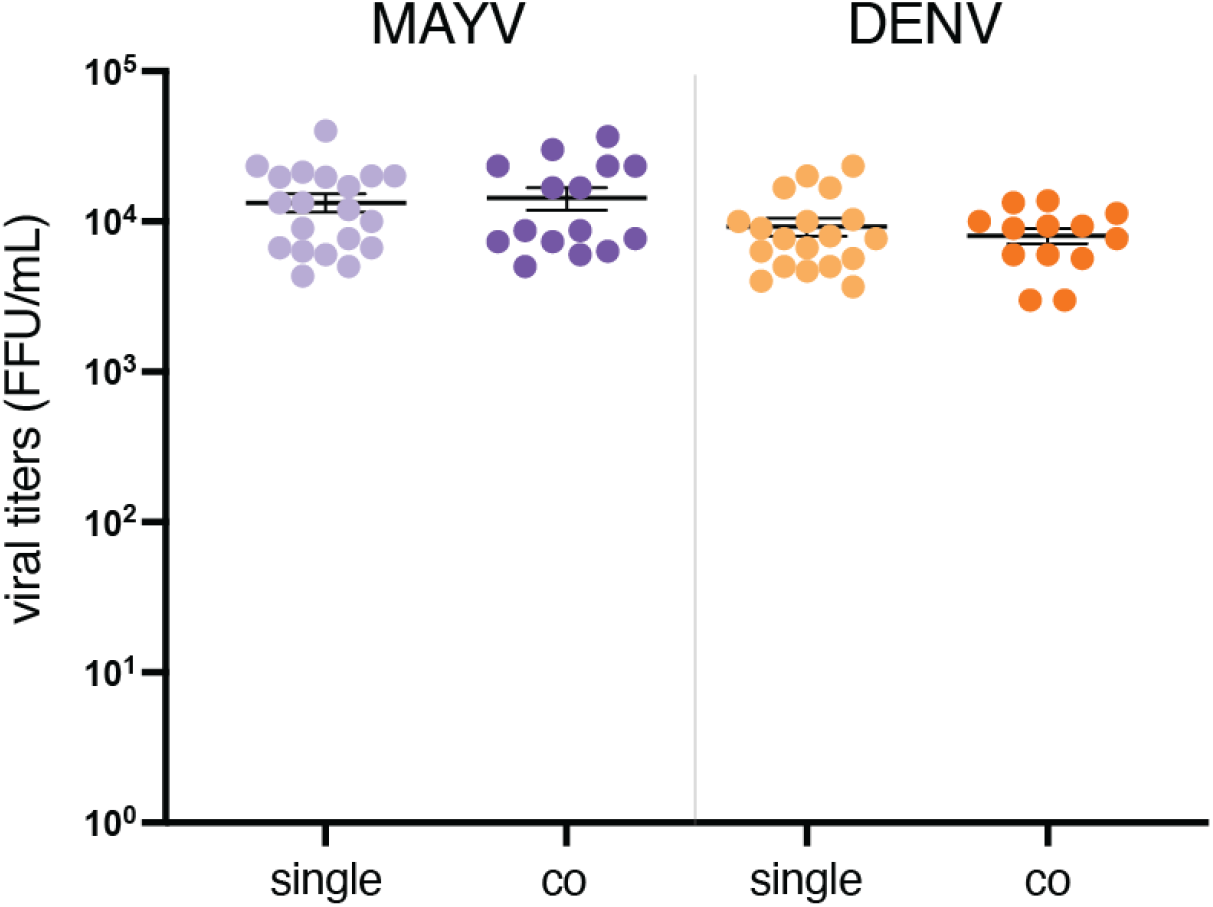
MAYV and DENV titers at initial infection (0dpi) correct?. Fully-engorged female mosquitoes were collected whole at the sorting step as a proxy to determine the infectious viral particles delivered to each mosquito. Each point represents a sample of an individual mosquito tissue. Purple shades depict MAYV titers and orange shades depict DENV. For co-infected mosquitoes, both viruses were tested on the same sample.

